# Galectin-8 Modulates Membrane CD44v Localization and Tempers STAT3 Signaling in Gastric Metaplasia

**DOI:** 10.64898/2026.06.07.729556

**Authors:** Xiaobo Lin, Xuemei Liu, Gabriel Nicolazzi, Yehiel Zick, Jeffrey W. Brown

## Abstract

Galectin-8 is a lectin that binds N-acetyllactosamine moieties with preference towards those with acidic, terminal modifications (-O-sialyated, 3’-O-sulfated). As CD44-variants (CD44v) are biomarkers of metaplasia and cancer, a hyaluronic acid receptor that modulates STAT3 signaling, and specifically expresses 3’-Sialyl-Le^A/X^ glycotopes, we asked whether galectin-8 might play a role in gastric metaplasia. Using a synchronous, chemically induced murine model that produces gastric spasmolytic polypeptide expressing metaplasia (SPEM), we compared *Lgals8*^−/–^ mice to congenic wild-type C57BL/6J mice. We found that galectin-8 was necessary for membrane localization of CD44v on SPEM cells at the base of the glands, suggesting a physical interaction between galectin-8 and CD44v. Metaplastic glands from *Lgals8*^*–/–*^ mice had an increase in nuclear pSTAT3 compared to C57BL/6J mice, suggesting that galectin-8 restrains CD44 -> STAT3 signaling. This effect was more prominent in the neck compared to the base, which has greater abundance of CD44v after injury in *Lgals8*^*–/–*^ mice. Derepression of STAT3 signaling may explain why low galectin-8 levels is associated with a worse prognosis in gastric cancer.

## INTRODUCTION

Galectins are a family of lectins that bind N-Acetyllactosamine sugars. At least three of these galectins (galectin-3, galectin-4, and galectin-8) preferentially associate with N-Acetyllactosamine that contains an acidic terminal moiety (-O-Sialyated, 3’-O-sulfated) 3’-site of galactose, an epitope that our group and others have shown is specifically expressed in high-risk metaplasia and cancer throughout the gastrointestinal foregut. Accordingly, specific roles for galectin-3(1) and galectin-4 have been reported in foregut cancer biology; however, a role of galectin-8 has not been similarly explored.

We have recently reported that galectin-8 is expressed and presumably secreted from gastric tuft, pit, proliferative, and endothelial cells of the murine stomach,(2) but that *Lgals8*^*–/–*^ mice do not have a significant phenotype at homeostasis when compared to age matched C57BL/6J mice.(2) Since CD44-variants have been reported to associate with galectin-8 in the setting of synovial inflammation(3) potentially through 3’-Sialylated lewis glycotopes that are specifically expressed on CD44v,(4) we asked whether galectin-8 alters CD44v signaling(5, 6) in the setting of gastric metaplasia.

## MATERIALS and METHODS

All experiments using animals followed protocols approved by the Washington University in St. Louis, School of Medicine Institutional Animal Care and Use Committee and conducted in accordance with the relevant guidelines and regulations. WT C57BL/6J mice were purchased from Jackson Laboratories (Bar Harbor, ME). All *Lgals8*^*–/–*^ mice used here were of the *Lgals8*^*–/–*^*/Mmrn1*^*WT*^*/Snca*^*WT*^ genotype.(2)

### Chemically-Induced SPEM

Tamoxifen powder (Toronto Research Chemicals) was initially solubilized in 100% ethanol via sonification after which it was emulsified in sunflower oil (Sigma-Aldrich) at a 9 Oil:1 EtOH ratio. Tamoxifen (5mg / 20 gm body weight) was injected intraperitoneally for 2 days and mice were euthanized for histologic examination at 48 or 72 hours after the first injection. All mouse experiments were performed on mice aged 6-10 weeks and both sexes were utilized indiscriminately as prior studies have demonstrated an identical phenotype. All injections and endpoints took place between 2 pm and 4 pm to avoid circadian effects.

### qRT-PCR

RNA was isolated using RNeasy (Qiagen) per the manufacturer’s protocol. The integrity of the mRNA was verified with a BioTek Take3 spectrophotometer and electrophoresis on a 2% agarose gel. RNA was treated with DNase I using in-column digestion (Qiagen), and 1 μg of RNA was reverse-transcribed with SuperScript III (Invitrogen) following the manufacture’s protocol. Measurements of cDNA abundance were performed by qRT– PCR using Applied Biosystems QuantStudio 3 Real-time PCR system. PowerUp^™^ SYBR^™^ Green Master Mix (Thermo Scientific) fluorescence was used to quantify the relative amplicon amounts of each gene. Primer sequences are located in the key resource table.

### Western Blot Analysis

Approximately 75 mg mouse corpus stomach was sonicated in RIPA buffer with 1× protease inhibitor cocktail (Thermo). Protein was separated using NuPAGE 10% Bis-Tris gels and transferred to an Invitrogen nitrocellulose membrane. Membranes were then blocked with 5% bovine serum albumin (BSA) in PBS and incubated overnight at 4°C with primary antibodies (see Reagent Table for antibodies). The membranes were rinsed in Tris buffered saline (TBS) pH 8.0, incubated 1 hour at room temperature in secondary antibody in 5% BSA in PBS. Beta-tubulin antibody was used as a control to ensure equal loading of protein in each gel lane. Membranes were imaged with Licor Odessey CLX and processed with associated ImageStudio software.

## RESULTS

We have previously reported that relative to congenic C57BL/6J, *Lgals8*^*–/–*^ mice do not have a significant phenotype at homeostasis.(2) In our chemically induced murine model of metaplasia, *Lgals8* levels did not significantly change during transition into spasmolytic polypeptide expressing metaplasia (SPEM) (Figure 1A). This result is likely due to *Lgals8* being absent from the cells that transform into SPEM, namely chief cells.(7) Similarly, we did not observe any defects in cathartocytosis, a newly described cellular process that facilitates rapidly downscaling of cellular machinery during the injury response (8) as evidenced by the presence of luminal material from the endoplasmic reticulum after injury (Supplemental Figure 1). However, when we investigated whether the absence of LGALS8 affected SPEM, we found that LGALS8 was necessary for outer membrane localization of CD44v on chief cells, but not elsewhere in the gland (Figure 1B,C).

**Figure 1.**
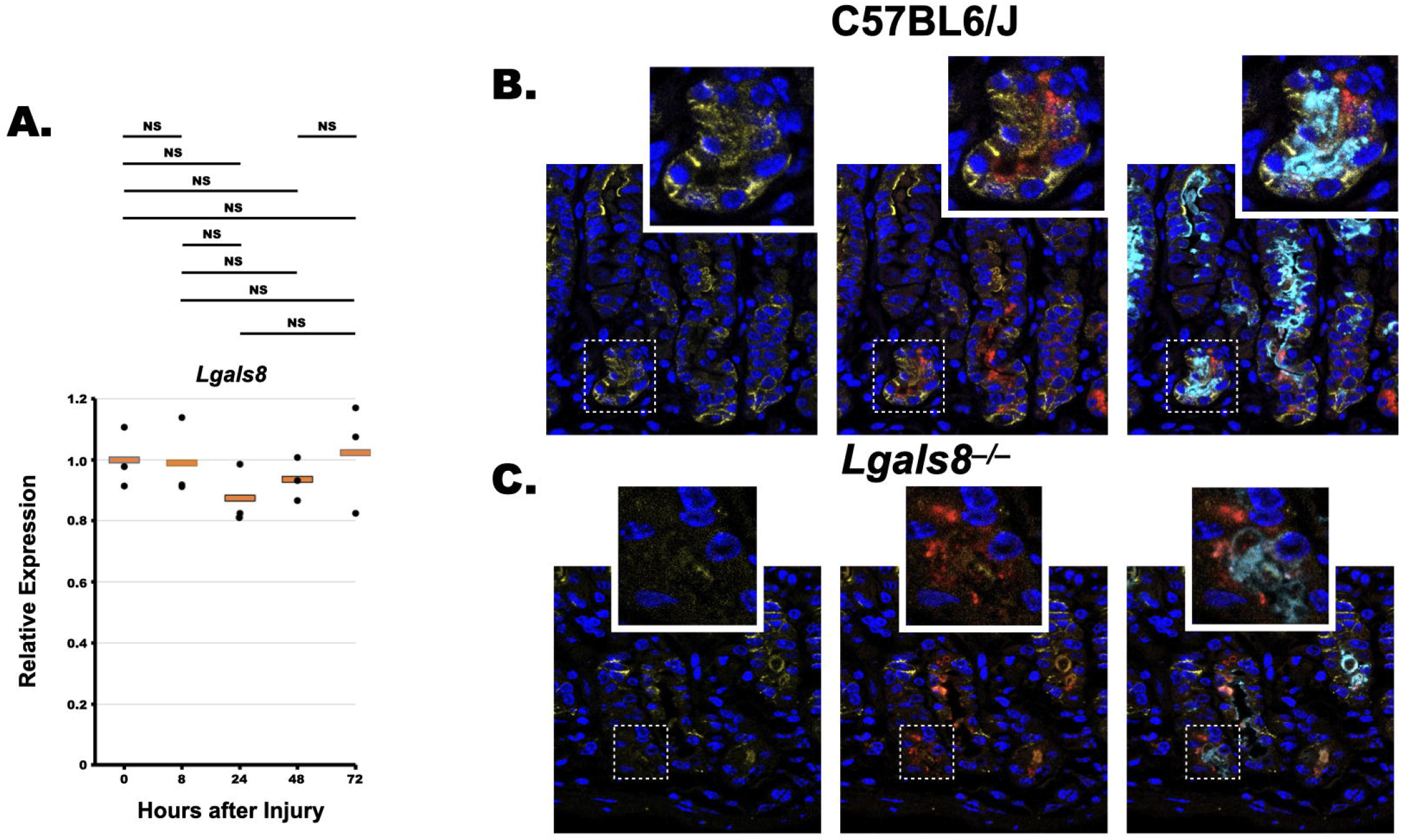
Galectin-8 stabilizes membrane CD44v in metaplasia. **A**. *Lgals8* levels are unaltered as the murine gastric glands transition to metaplasia. **B**. Confocal microscopy of wild-type C57BL/6J gastric glands 48 hours after tamoxifen injection demonstrates membrane expression of CD44v in SPEM cells. **C**. Confocal microscopy of *Lgals8*^*–/–*^ gastric glands 48 hours after tamoxifen injection demonstrates minimal membrane CD44v in SPEM at the base of the gland. Note persistent membrane CD44v expression in the neck. CD44v (yellow), GIF (red), and GSII lectin (Cyan).

CD44v modulates downstream STAT3 signaling in gastric metaplasia.(6) Accordingly, we asked whether STAT3 signaling was altered in the *Lgals8*^*–/–*^ mice. We found that nuclear pSTAT3 was significantly increased in *Lgals8*^*–/–*^ mice, relative C57BL/6J mice (Figure 2A,B). This effect was more pronounced in the neck region, which expressed greater membrane staining of CD44v in *Lgals8*^*–/–*^ mice (Figure 1), relative to the base of the gland. This histological result was confirmed with western blot analysis of nuclear fractions from *Lgals8*^*–/–*^ and C57BL/6J mice (Figure 2C,D).

**Figure 2.**
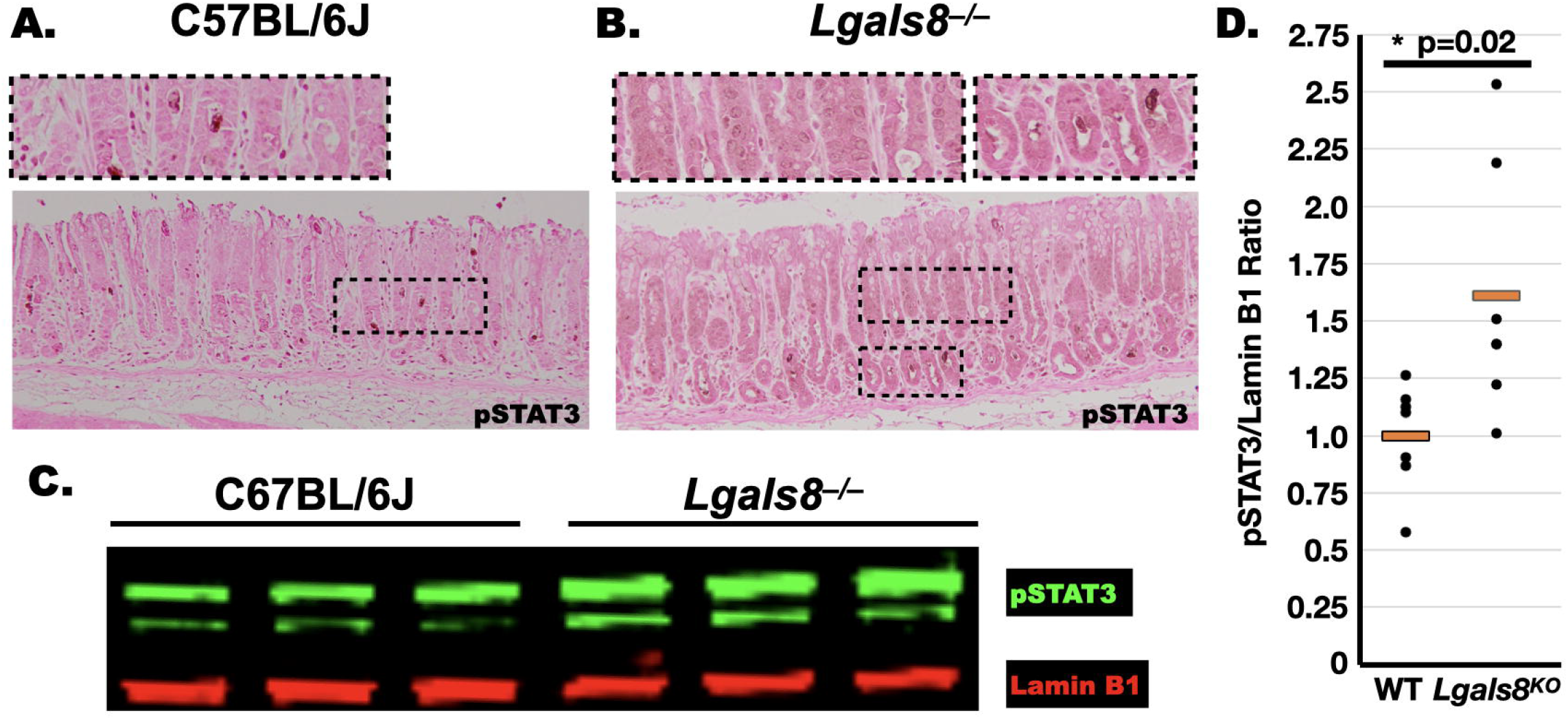
Galectin-8 tempers nuclear p-STAT3. **A**. p-STAT3 immunohistochemistry of C57BL/6J mice 48 hours after tamoxifen administration demonstrates reactivity only to cellular debris secreted via cathartocytosis. **B**. p-STAT3 immunohistochemistry of *Lgals8*^−/–^ mice 48 hours after tamoxifen administration demonstrates nuclear reactivity throughout the corpus gland. **C**. Western blot analysis of nuclear fractions from three C57BL/6J and three *Lgals8*^*–/–*^ mice demonstrates increased p-STAT3 when normalized to laminin. Quantification of normalized pSTAT3/laminin ratio. Dots represent individual mice, orange line is the average ratio. P-value is calculated with student T-test.

## DISCUSSION

Galectins are known to play numerous and diverse roles in metaplasia and cancer. Here we found that LGALS8 was necessary for outer membrane localization of CD44v in metaplastic SPEM cells at the base of the gland, but not in the neck (Figure 1B,C). This observation may result from divergent, cell-type specific glycotopes displayed on CD44v (and elsewhere in the neighboring extracellular space) or a different predisposition to endocytose CD44v upon extracellular binding / signaling. Accordingly, we observed that the increase in STAT3 signaling was more pronounced in the neck region compared to the base of the gland, which is consistent with a greater abundance of CD44v in the neck cells relative to the base in *Lgals8*^*–/–*^ mice.

Increased STAT3 signaling via CD44 promotes pro-cancer phenotypes including cell survival, proliferation, invasion, and migration.(9) Restraining aggressive phenotype is of upmost importance in paligenotic cellular transformations as these processes involve licensing long-lived differentiated cells that have banked potentially oncogenic mutations back into the cell cycle. Our data suggests that LGALS8 would restrain such aggressive phenotypes by decreasing STAT3 signaling through CD44 and may serve to explain why low galectin-8 protein levels are correlated with poor survival in gastric cancer.(10)

This study is not without limitations. The chemically-induced injury model utilized here generates murine SPEM that is both transcriptionally and histologically highly similar to that generated by *Helicobacter* infection; however, neither of these models produce gastric cancer. Future studies in pro-oncogenic genetic background is necessary to determine whether the increased nuclear pSTAT3 in *Lgals8* null mice results in an increased predilection to develop dysplasia and cancer.

**Supplementary Figure 1.**
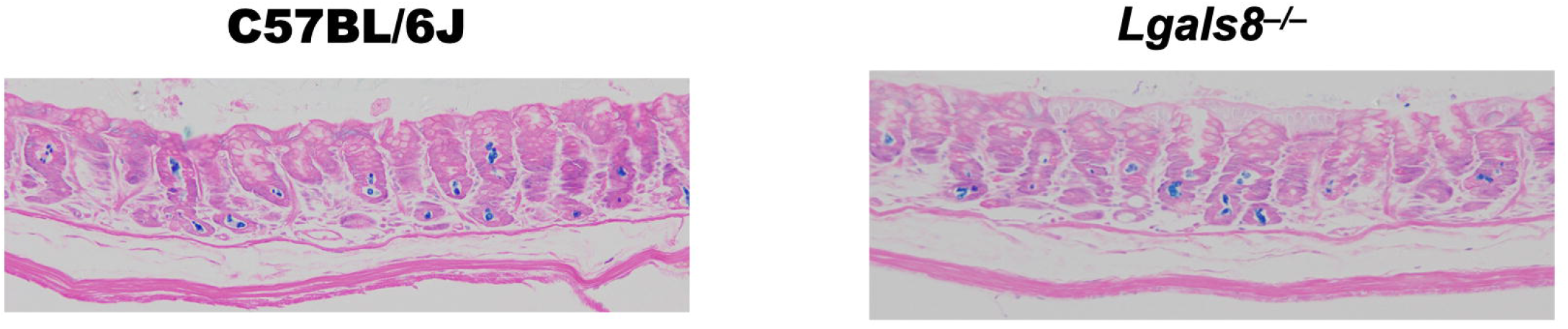
Demonstration of Cathartocytosis in *Lgals8*^*–/–*^ mice. Azure A staining (with Eosin counterstain) demonstrates the expulsion of endoplasmic reticulum contents 48 hours after injury in both (**A**) Wild-type C57BL/6J and (**B**) *Lgals8*^*–/–*^ mice.

## KEY RESOURCE TABLE

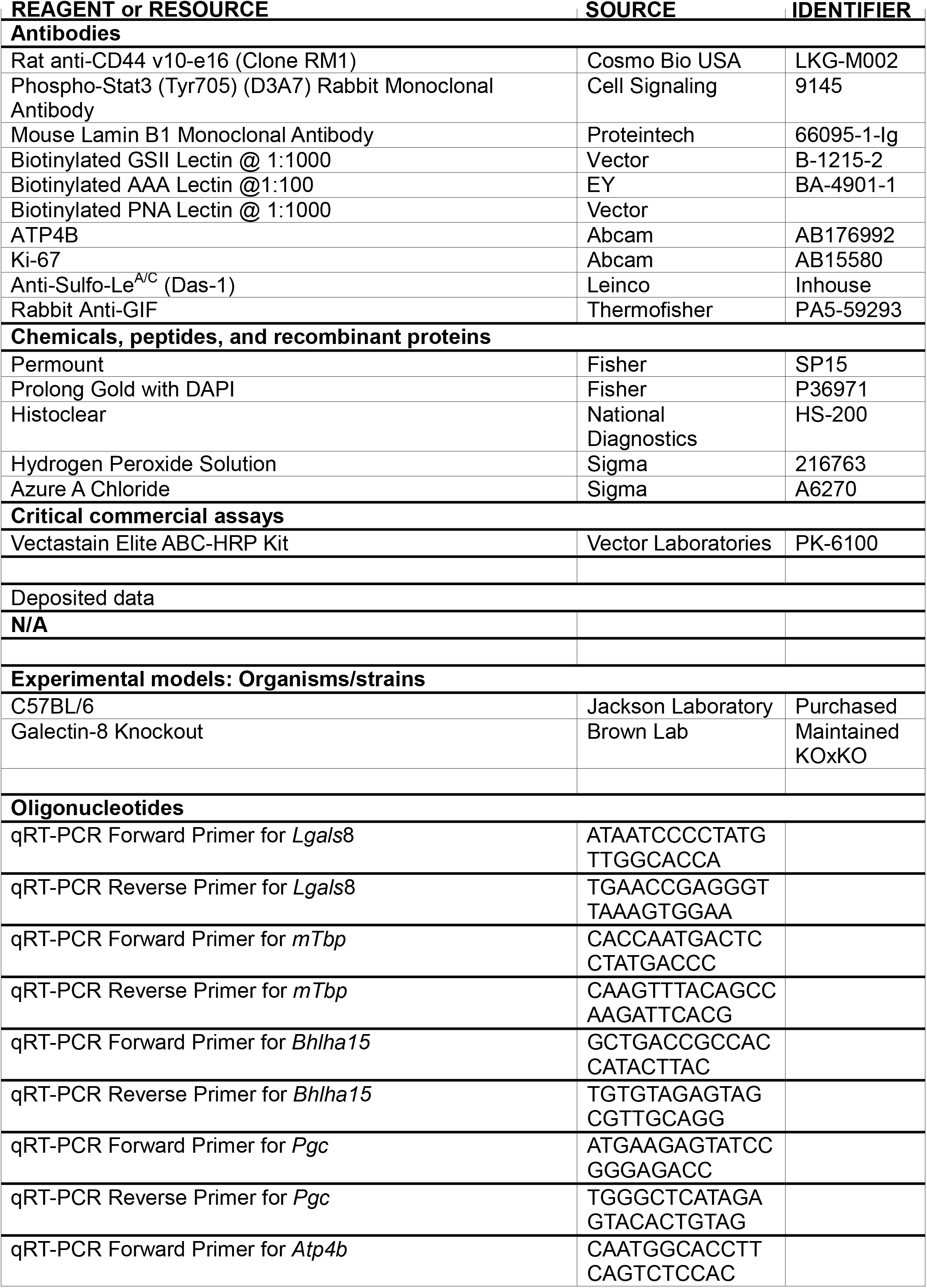
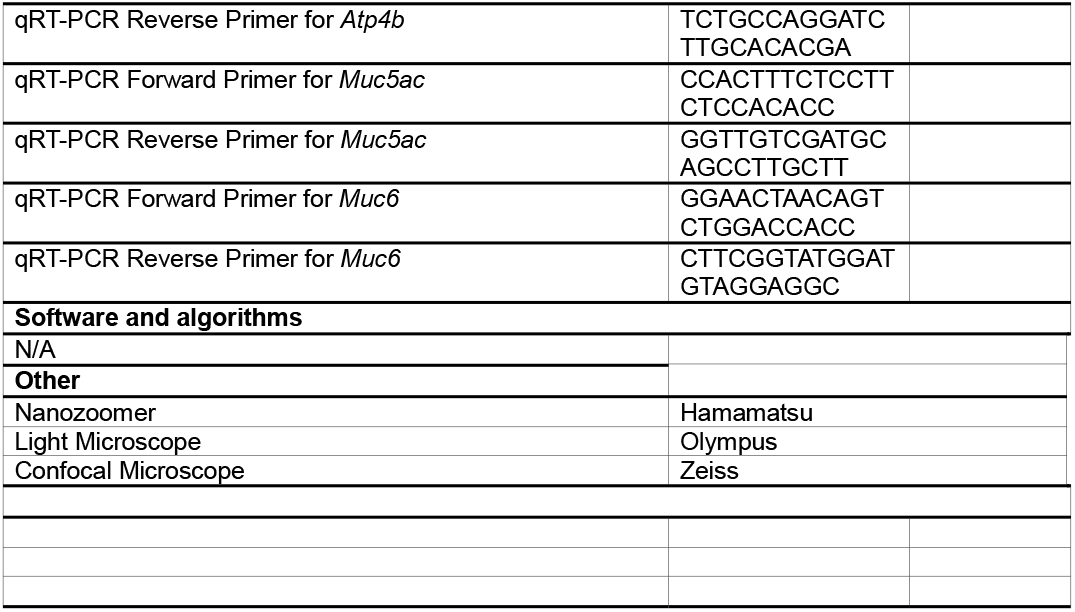

